# Alternative splicing variants of Arabidopsis G protein β subunit AGB1 function in plant development and endoplasmic reticulum stress response

**DOI:** 10.1101/2023.11.26.568767

**Authors:** Yueh Cho

**Affiliations:** Institute of Plant and Microbial Biology, Academia Sinica, Taipei, 115201, Taiwan

**Keywords:** AGB1, alternative splicing, development, endoplasmic reticulum, ER stress, Gβ protein, Heterotrimeric G protein

## Abstract

The Arabidopsis heterotrimeric G protein β subunit, GTP BINDING PROTEIN BETA 1 (AGB1), has multiple functions in plant development and response to various environmental stimuli including endoplasmic reticulum (ER) stress. However, how the single gene produces the pleiotropic effect remains elusive. Here, we show that AGB1 has 4 alternative splice isoforms with isoform-specific features. AGB1.2 failed to rescue the *agb1-3* mutant defects and thus was considered a non-functional isoform. Although AGB1.1 and AGB1.4 were both localized at the plasma membrane and the ER, AGB1.1 fully rescued the defects of *agb1-3,* and AGB1.4 only partially rescued the defects even though its transcript level was higher than that of AGB1.1. Intriguingly, AGB1.3 was localized at the nucleus and further enhanced the leaf-shape phenotype of *agb1-3*. The protein structure of AGB1.3 is unique because of the termination of translation in the 7th-WD40 motif by alternative splicing, which produced an incomplete propeller structure. AGB1.1 and AGB1.4 but not AGB1.3 interacted with the Gγ subunits, AGG1, AGG2, and AGG3, possibly because of the lack of the 7th-WD40 motif in AGB1.3. AGB1 may produce its multifaceted functions in plant development and ER stress tolerance via its alternative splice isoforms with distinct structural features and subcellular localization.

## INTRODUCTION

The heterotrimeric guanine nucleotide-binding protein (G protein) consists of ⍺, β and γ subunits that serve as a signal mediator coupled with the plasma membrane (PM)-spanning G-protein-coupled receptors (GPCRs) and effectors (Marinissen and Gutkind, 2001). G⍺ forms a trimeric complex upon external stimuli with the Gβγ heterodimer for interaction with downstream effectors (Lee and Assmann, 1999). The *Arabidopsis thaliana* genome encodes 4 G⍺ subunits [one canonical *GPA1* subunit (Ma et al., 1990) and 3 non-canonical G⍺ subunits, *XLG1* (Lee and Assmann, 1999), *XLG2* (Ding et al., 2008, Zhu et al., 2009) and *XLG3* (Ding et al., 2008, Chakravorty et al., 2015)], one Gβ subunit [*AGB1*(Weiss et al., 1994)] and 3 Gγ subunits [*AGG1* (Mason and Botella, 2000), *AGG2* (Mason and Botella, 2001) and *AGG3* (Chakravorty et al., 2011)]. Loss-of-function mutations of *AGB1* exhibit strikingly diverse developmental defects such as compact and rounded leaves (Lease et al., 2001), short sepals (Ullah et al., 2003), short silique with blunt tip (Lease et al., 2001) and large biomass (Ullah et al., 2003). In addition to the developmental defects, *AGB1* mutations alter the sensitivity to a broad range of environmental stresses including salinity stress (Colaneri et al., 2014), drought stress (Xu et al., 2015), ozone (O3)-triggered oxidative stress (Joo et al., 2005), pathogen attack (Cheng et al., 2015) and endoplasmic reticulum (ER) stress (Chen and Brandizzi, 2012, Cho et al., 2015). Because of its importance in plant growth and stress tolerance, AGB1 has been extensively investigated for its molecular functions. However, how the single *AGB1* gene as a responsible β subunit can have diverse roles in plants remains elusive.

Like animal cells, in plants, the interaction with Gγ subunits is important for AGB1 function because knocking out all 3 Gγ subunits, AGG1, AGG2, and AGG3, caused a phenocopy of the *agb1* mutants (Thung et al., 2012). Indeed, AGB1 interacts with Gγ subunits in Arabidopsis: yeast-based heterologous assays showed the interaction of AGB1 with AGG1 (Mason and Botella, 2000), AGG2 (Klopffleisch et al., 2011), and AGG3 (Wolfenstetter et al., 2015). AGB1 is also a member of the tryptophan, aspartic acid (WD)-repeat family proteins that are involved in cellular signaling pathways because of a role in scaffolding protein complexes that bind multiple partner proteins. In fact, approximately 96 binding proteins of AGB1 are reported/predicted in Arabidopsis (BioGRID, https://thebiogrid.org). AGB1 is assumed to be localized at the PM because of the function of the heterotrimeric G protein complex. Cumulative evidence has localized AGB1 at the PM (Wang et al., 2007), AGB1 was also suggested to be localized at the ER and nucleus (Anderson and Botella, 2007, Wang et al., 2007). AGB1 in the nucleus interacts with B-box (BBX)-containing zinc-finger transcription factor (BBX21) for hypocotyl elongation (Xu et al., 2017), brassinosteroids transcription factor BRI1-EMS-SUPPRESSOR 1 (BES1) for cell division (Zhang et al., 2017), blue light photoreceptor CRYPTOCHROME 1 (CRY1) - basic leucine zipper transcriptional factor ELONGATED HYPOCOTYL 5 (HY5) (Lian et al., 2018), red light photoreceptor PHYTOCHROME B (phyB) - basic helix-loop-helix transcriptional factor PHYTOCHROME-INTERACTING FACTOR 3 (PIF3) (Xu et al., 2019) for photomorphogenesis, and MITOGEN-ACTIVATED PROTEIN KINASE 6 (MPK6) (Xu et al., 2015) for drought tolerance. Hence, AGB1 is a multifunctional protein in Arabidopsis, but how this is achieved by a single gene remains unanswered.

Alternative splicing of precursor mRNAs from multiexon genes increases the diversity of their potential gene expression profiles. In plants, almost 90% of the protein-coding genes possess introns, and the extent of alternative splicing has been reported to range from 40% to 60% of intron-containing genes (Marquez et al., 2012, Staiger and Brown, 2013). Stress-responsive genes in plants often use alternative splicing to maintain the balance of active and non-active isoforms under stress conditions. The heat shock transcription factor A2 (*HsfA2*) is a case for temperature-induced alternative splicing in Arabidopsis (Liu et al., 2013). *HsfA2-III*, an alternative splice isoform, produces a small truncated protein at high temperature, which is localized to the nucleus and binds the promoter of *HsfA2* to control *HsfA2* expression (Liu et al., 2013). Under cold conditions, alternative splicing of the Arabidopsis *INDERMINATE DOMAIN 14* (*IDD14*) regulates starch metabolism by producing 2 splice isoforms that encode the functional transcription factor and truncated form lacking one of the zinc finger motifs (Seo et al., 2011). In rice, alternative splicing of *Oryza sativa DEHYDRATION-RESPONSIVE ELEMENT-BINDING PROTEIN2* (*OsDREB2B*) is required for rapid production of the functional OsDREB2B protein in response to abiotic stress (Matsukura et al., 2010). AGB1 transcripts consist of 4 alternatively spliced isoforms, *AGB1.1* to *AGB1.4*, based on TAIR (https://www.arabidopsis.org). Thus, the multifaceted function of AGB1 may be achieved by alternative splicing.

In this study, we demonstrate that *AGB1* produces 4 alternative splice isoforms with isoform-specific features. AGB1.1 and AGB1.4 were both localized at the PM and ER; AGB1.1 fully but AGB1.4 only partially rescued the defects of the *agb1-3* mutant. However, the transcript level was higher for *AGB1.4* than *AGB1.1*, which suggests a post-translational modification regulating both isoforms. AGB1.2 failed to rescue the defects of the *agb1-3* mutant, so it may be a non-functional isoform. Intriguingly, AGB1.3 was localized at the nucleus and its expression further enhanced the leaf shape phenotype of the *agb1-3* mutant. The protein structure of AGB1.3 is unique because of the termination of translation in the 7th-WD40 motif by alternative splicing that produces an incomplete propeller structure. AGB1.1 and AGB1.4 but not AGB1.3 interacted with the Gγ subunits AGG1, AGG2, and AGG3, possibly because of lack of the 7th-WD40 motif in AGB1.3. AGB1 may produce its multifaceted functions in plant development and ER stress tolerance via its alternative splice isoforms with distinct structural features and subcellular localization.

## RESULTS

### *ProAGB1:AGB1-Ven agb1-3* plants complement the phenotypes of the *agb1-3* mutant

To address how the single gene, *AGB1*, fulfills multiple roles with diverse subcellular localizations in Arabidopsis, we first created transgenic plants harboring a genomic sequence of *AGB1* fused C-terminally to the Venus (Ven) fluorescent protein in an *agb1-3* (SALK_061896) genetic background (*ProAGB1:AGB1-Ven agb1-3*, hereafter called AGB1-VEN) (Figure 1A). The *agb1-3* is a null mutant of *AGB1* (Chen and Brandizzi, 2012). In the T2 generation, we isolated 24 independent transgenic lines with the transgene and selected 3 representative lines (lines #6, #10 and #17). For genomic complementation assay, the T3 generation was used (lines #6-3, #10-1, and #17-2). Consistent with previous studies (Lease et al., 2001, Ullah et al., 2003), *agb1-3* showed diverse developmental defects such as compact and rounded rosette leaves (Figure 1B, 1C), short sepal (Figure 1D), and short silique with blunt tip (Figure 1E, 1F, 1G). These developmental defects were complemented in AGB1-VEN plants (Figure 1B to 1G). We next tested whether AGB1-VEN was also able to rescue the *agb1-3* defective ER stress sensitivity (Chen and Brandizzi, 2012). When we grew wild-type, *agb1-3* and AGB1-VEN plants on MS-agar plates for 2 weeks, *agb1-3* grew better than the wild type because *agb1-3* is known to show increased biomass of seedlings (Figure 1H and 1I) (Lease et al., 2001, Ullah et al., 2003, Chen et al., 2006). In contrast, in the presence of the ER stress inducer tunicamycin (TM), fresh weight was significantly less for *agb1-3* than the wild-type and AGB1-VEN plants (Figure 1H and 1I). Likewise, the other 2 transgenic lines of AGB1-VEN complemented the ER-stress phenotype of *agb1-3* (Supplemental Figure S1). Therefore, AGB1-VEN was functional in both plant development and ER stress tolerance.

**Fig. 1.**
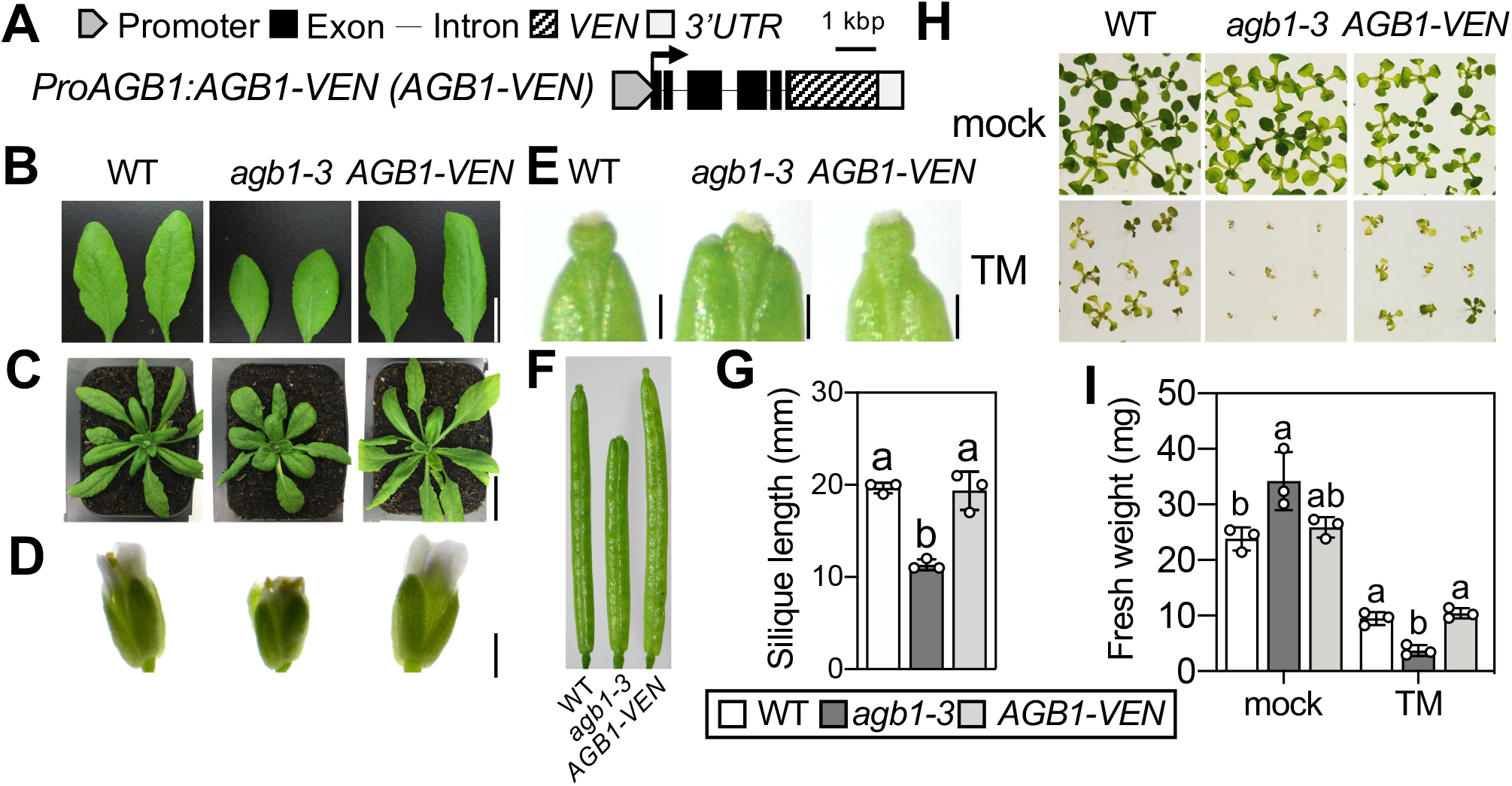
Functional complementation of hypersensitivity to ER stress in *agb1-3* mutant. (A) The gene structure of transgenic complementation lines (ProAGB1:AGB1-Ven *agb1-3*, AGB1-VEN). Morphological observation of 18-day-old rosette leaves (B), 24-day-old plants (C), 35-day-old flowers (D) and 42-day-old siliques (F) from wild-type (WT), *agb1-3* mutant (*agb1-3*) and transgenic complementation line (AGB1-VEN). Images of silique tip were shown in (E) and silique length was quantified from 16 of mature plants shown by mean ± SD (G). Representative images (H) and fresh weight (I) of 14-day-old WT, *agb1-3* mutant and AGB1-VEN plants grown on 1/2 MS containing dimethyl sulfoxide (mock) or tunicamycin (TM). Fresh weight of the seedlings in H was measured individually (n=25), and shown as mean ± SD (mg) in I. Different letters indicate significant difference at P < 0.05 as determined by one-way ANOVA. Values with the same letter are not significantly different. Three biologically independent experiments were performed with similar results. Scale bars; 1 cm in (B), 2 cm in (C), 1 mm in (D) and 3 mm in (E).

### AGB1 is located at the PM and ER *in vivo*

To observe the functional AGB1 protein *in vivo*, we observed the subcellular localization of AGB1 in AGB1-VEN plants (line #6-3). AGB1-VEN in the root epidermis of 7-day-old plants was detected at the cell boundary in addition to the network structure (Figure 2A and 2E), which overlapped with the PM marker FM4-64 (Figure 2A, 2B, 2C) and ER tracker (Figure 2E, 2F, 2G), respectively. Similar results were observed in the other representative lines, #10-1 and #17-2 (Supplementary Figure S2). An early study using Arabidopsis wild-type plants expressing cDNA of AGB1 fused to GFP driven by the 35S promoter reported AGB1-GFP localized to the PM and in the nucleus in leaf epidermal cells (Anderson and Botella, 2007); however, we found PM but not nuclear localization. To validate the ER localization of AGB1, we further performed a co-transient assay with AGB1-VEN (Figure 2I) and the ER-resident protein mRFP-HDEL (Figure 2J) co-expressed in Arabidopsis wild-type protoplasts. As the yellow signals show in the merged image (Figure 2K), AGB1-VEN co-localized with mRFP-HDEL but not chlorophyll autofluorescence (Figure 2L). Thus, AGB1-VEN localized in the PM and ER in stable transgenic plants.

**Fig. 2.**
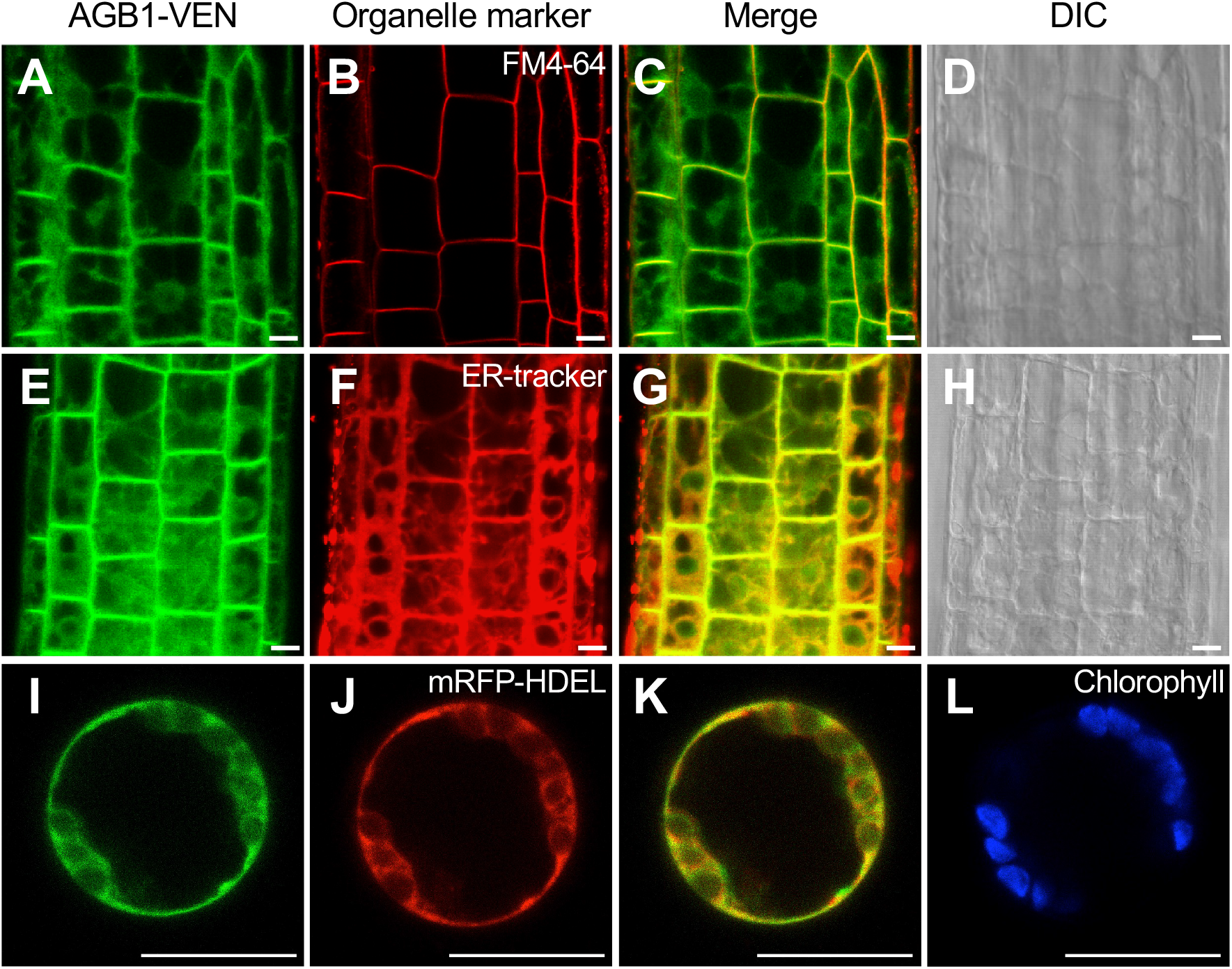
Subcellular localization of AGB1. Representative images AGB1-VEN in ProAGB1:AGB1-Ven *agb1-3* plants (A-H) or in wild-type protoplasts transiently expressing ProAGB1:AGB1-Ven (I-L). Fluorescence of AGB1-VEN (Green, A, E) and staining of the plasma membrane by FM4-64 (Red, B) or the ER by ER-tracker (Red, F). Cell structure is shown in differential interference contrast (DIC) image at the right (D, H) in root of 7-day-old ProAGB1:AGB1-Ven *agb1-3* plant. The merged images are shown, respectively (C, G). AGB1-VEN (Green, I) was co-transduced with the ER-marker (Red, mRFP-HDEL, J) into the wild-type protoplast. The merged image of AGB1-VEN and the ER-maker is shown (K). The chlorophyll autofluorescence is shown on the right in blue signals (L). Scale bars, 10 µm.

### AGB1 encodes 4 alternative splice isoforms

According to the Arabidopsis Information Resource (TAIR, https://www.arabidopsis.org), genomic *AGB1* produces 4 alternative splice isoforms, *AGB1.1* to *AGB1.4*. *AGB1.1* contains full-length mRNA (1,850 bp), whereas *AGB1.2* (1,828 bp) lacks an N-terminal region corresponding to the first and second exons (Figure 3A and B). *AGB1.3* (1,642 bp) is produced from an alternative 3’ splice site in the 4th intron, which creates a stop codon in exon 5 (Figure 3A). Therefore, AGB1.3 lacks 30 amino acid residues at the C-terminus (Figure 3B) as compared with AGB1.1. *AGB1.4* (1,616 bp) is derived from an alternative 3’ splice site in the first intron, which causes a 15-nt deletion in the second exon (Figure 3A). Thus, the amino acid sequence of AGB1.4 is identical to that of AGB1.1 except for the deletion of 5 amino acid residues, VARYS, in the second exon (Figure 3A and 3B). Importantly, our functional transgenic line, AGB1-VEN, produces only 3 splice isoforms as a Ven fusion protein, AGB1.1-VEN, AGB1.2-VEN, and AGB1.4-VEN, but not for AGB1.3. Thus, the information for subcellular localization of AGB1.3 is missing in the observations shown in Figure 2. AGB1 has been known to form a beta-propeller protein structure by 7 repeatable WD40 domains that serves as a scaffold protein (Ullah et al., 2003) in addition to an alpha-helical coiled-coil domain at the N-terminus (Ullah et al., 2003). AGB1.2 and AGB1.4 contain incomplete coiled-coil domains, whereas AGB1.3 lacks the 7th WD domain. Because these differences will affect tertiary structures of protein as predicted by consulting the SWISS-MODEL web portal (https://swissmodel.expay.org) (Figure 3C), each splice isoform may function differentially *in vivo*.

**Fig. 3.**
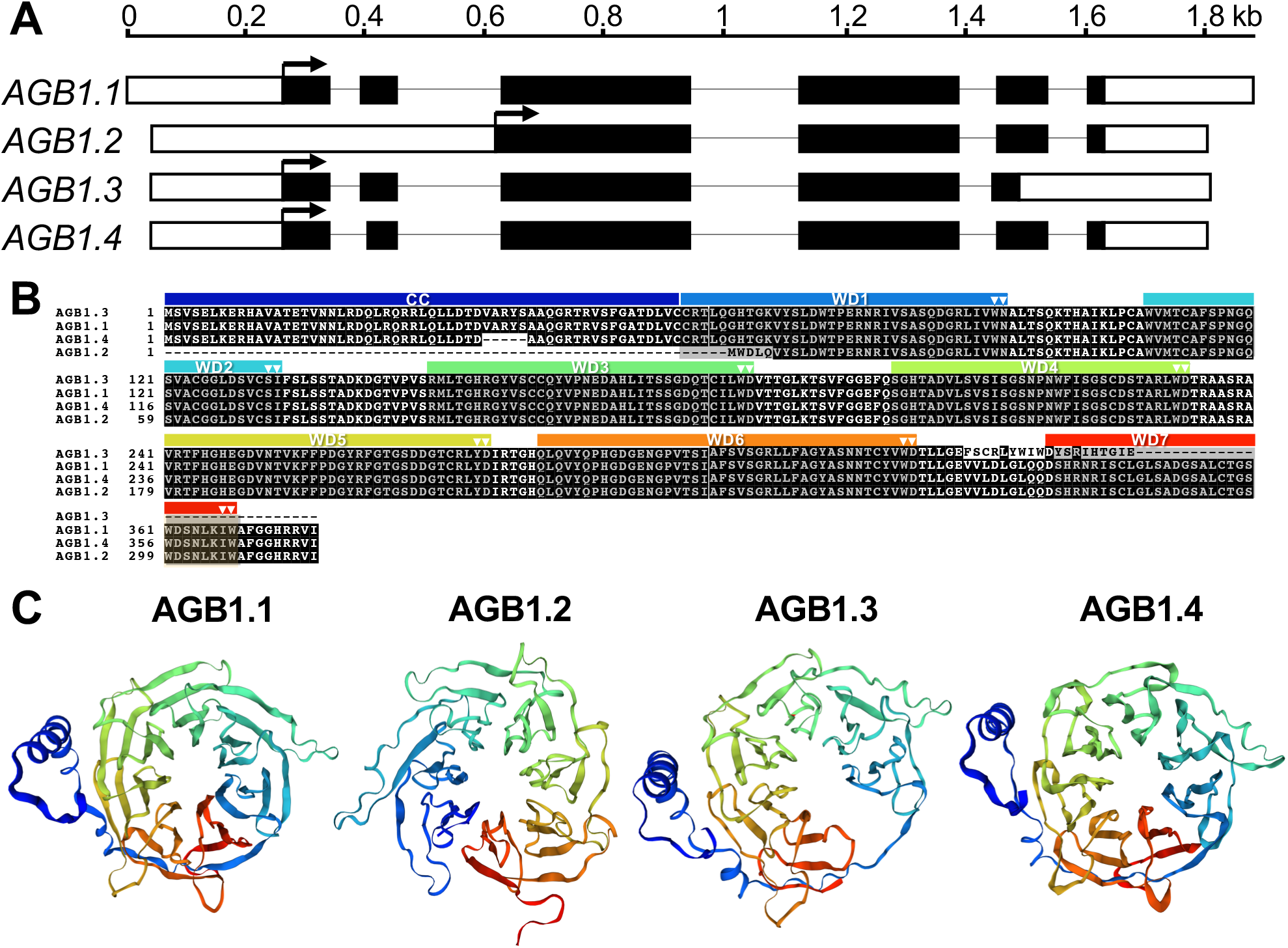
Schematic representation of 4 AGB1 splicing variants. (A) Schematic representation of the 4 splicing variants predicted for AGB1 (At4g34460). Black boxes represent exons, white boxes untranslated region, and arrows the position for start codon. (B) Amino acid alignment of the deduced AGB1 variant-coding sequences. The double triangles indicate the dipeptide for tryptophan-aspartic acid 40 (WD40) motif that is shaded with gray box. The dark blue box indicates the coiled-coil region (CC) and blue to red boxes the WD40 regions 1 to 7 (C) Tertiary structure prediction of 4 AGB1 variants from the SWISS-MODEL web portal (http://swissmodel.expasy.org).

### The 4 splice isoforms of AGB1 express differentially among organs and tissues

To investigate the tissue-specific expression of the 4 *AGB1* splice isoforms, we used RT-qPCR with different stages and tissues of wild-type plants (Figure 4). The transcript level of *AGB1.4* was always highest among the 4 splice isoforms in all tissues examined, but *AGB1.1* was also highly expressed next to *AGB1.4*. In contrast to *AGB1.1* and *AGB1.4*, the remaining 2 isoforms were rarely expressed in all tissues except *AGB1.2* in root and shoot of 14-day-old plants (Figure 4C and 4D). Indeed, *AGB1.3* was nearly undetectable in all tissues and organs examined. Thus, *AGB1.1* and *AGB1.4* may be the primary isoforms with respect to transcript level.

**Fig. 4.**
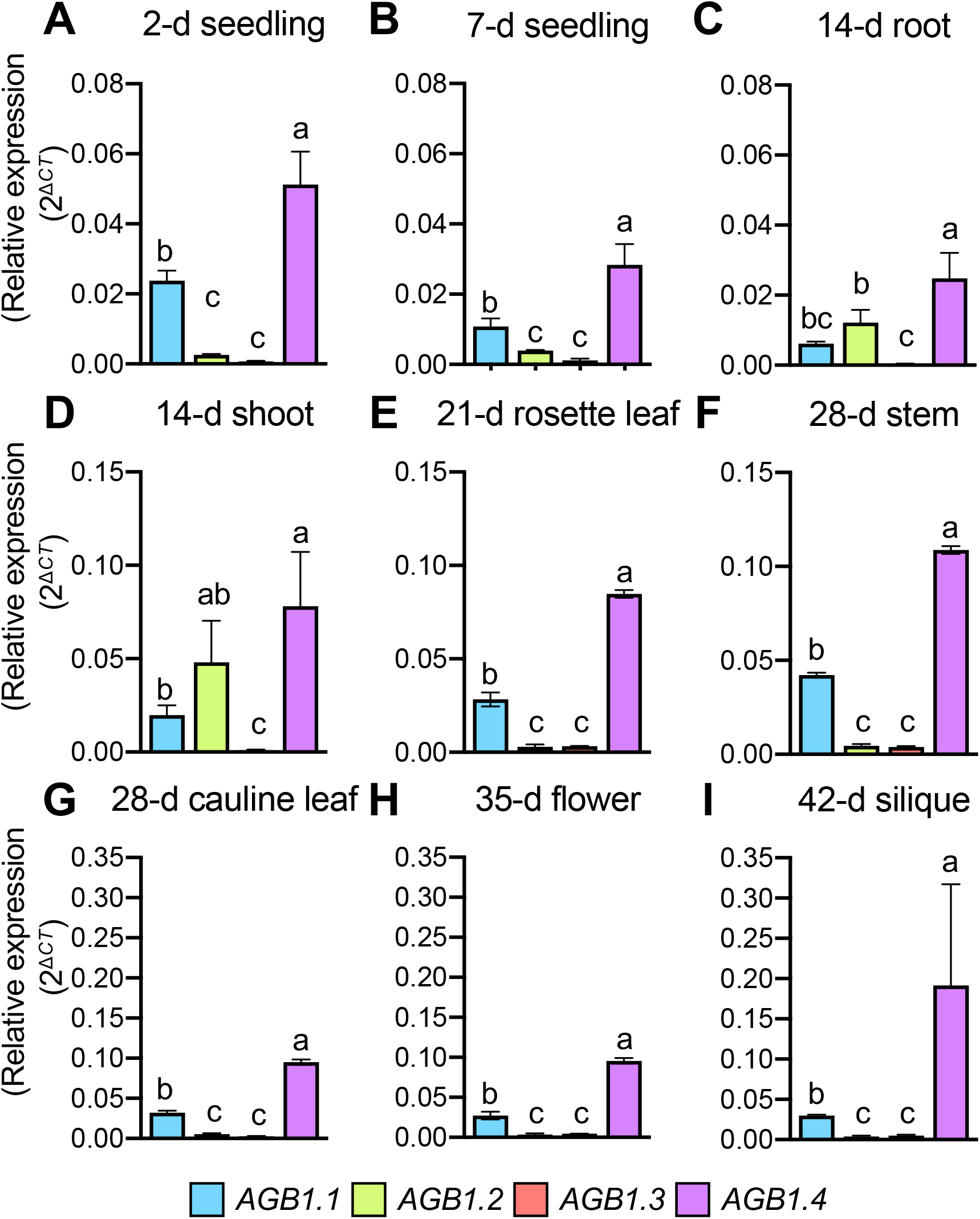
Expression profiles of 4 AGB1 variants. Transcript level of 4 AGB1 variants in different organs at different developmental stages in the wild type. RT-qPCR analysis of the expression of the AGB1 variants (AGB1.1, sky blue; AGB1.2, light green; AGB1.3, orange red; AGB1.4, light purple) in (A) 2-d seedling, (B) 7-d seedling, (C) 14-d root, (D) 14-d shoot, (E) 21-d rosette leaf, (F) 28-d stem, (G) 28-d cauline leaf, (H) 35-d flower (I) 42-d silique. Transcript levels were normalized to ACTIN2 and were shown as 2^ΔCT^. Data are mean ± SD from three biological replicates with three technical replicates in the same run. Different letters indicate significant difference at P < 0.05 as determined by one-way ANOVA. Values with the same letter are not significantly different.

### The splice isoforms of AGB1 differentially complement defects in plant development

To dissect the protein functions in distinct splice isoforms, we further created stable transgenic plants expressing coding sequences of each *AGB1* isoform fused with the VEN fluorescent reporter driven by the endogenous promoter in an *agb1-3* mutant background (*ProAGB1: AGB1.x-Ven agb1-3*, hereafter called AGB1.x-V). In the T2 generation, we isolated at least 48 independent transgenic lines with the respective transgenes and selected 3 representative lines for each construct. We first grew AGB1.1-V, AGB1.2-V, AGB1.3-V and AGB1.4-V with WT, *agb1-3*, and AGB1-VEN on soil for 24 days (Figure 5A). The phenotype of leaves in *agb1-3* was fully rescued by AGB1-VEN (the top-right image in Figure 5A), which is consistent with previous results (Figure 1B, 1C), and 3 representative lines of AGB1.1-V (lines #12-4, #18-7 and #22-7) also complemented these phenotypes (Figure 5A). AGB1.4-V (lines #16-3, #18-4 and #20-1) partially complemented the phenotype, but AGB1.2-V (lines #20-8, #22-2, #24-4) fail to do so (Figure 5A). Because of more rounded leaves, AGB1.3-V showed an enhanced phenotype as compared with the *agb1-3* mutant in all 3 transgenic plants (lines #1-7, #5-8 and #9-5). We further observed siliques because *agb1-3* plants produce short siliques with characteristic flat tips (Figure 1E) (Lease et al., 2001, Ullah et al., 2003, Chen et al., 2006). Consistent with the observation of leaves (Figure 5A), AGB1.1-V fully complemented the phenotype of both flat tips and short siliques (Figure 5B and 5C). Indeed, the WT and 3 AGB1.1-V lines (lines #12-4, #18-7 and #22-7) did not differ in silique length (Figure 5D). Similar to the results obtained in leaves (Figure 5A), AGB1.4-V partially complemented the silique phenotypes, but AGB1.2-V and AGB1.3-V failed to do so (Figure 5B, C, D). Hence, AGB1.1-V and AGB1.4-V complemented the phenotypes of *agb1-3* fully and partially, respectively, but neither AGB1.2-V nor AGB1.3-V complemented them.

**Fig. 5.**
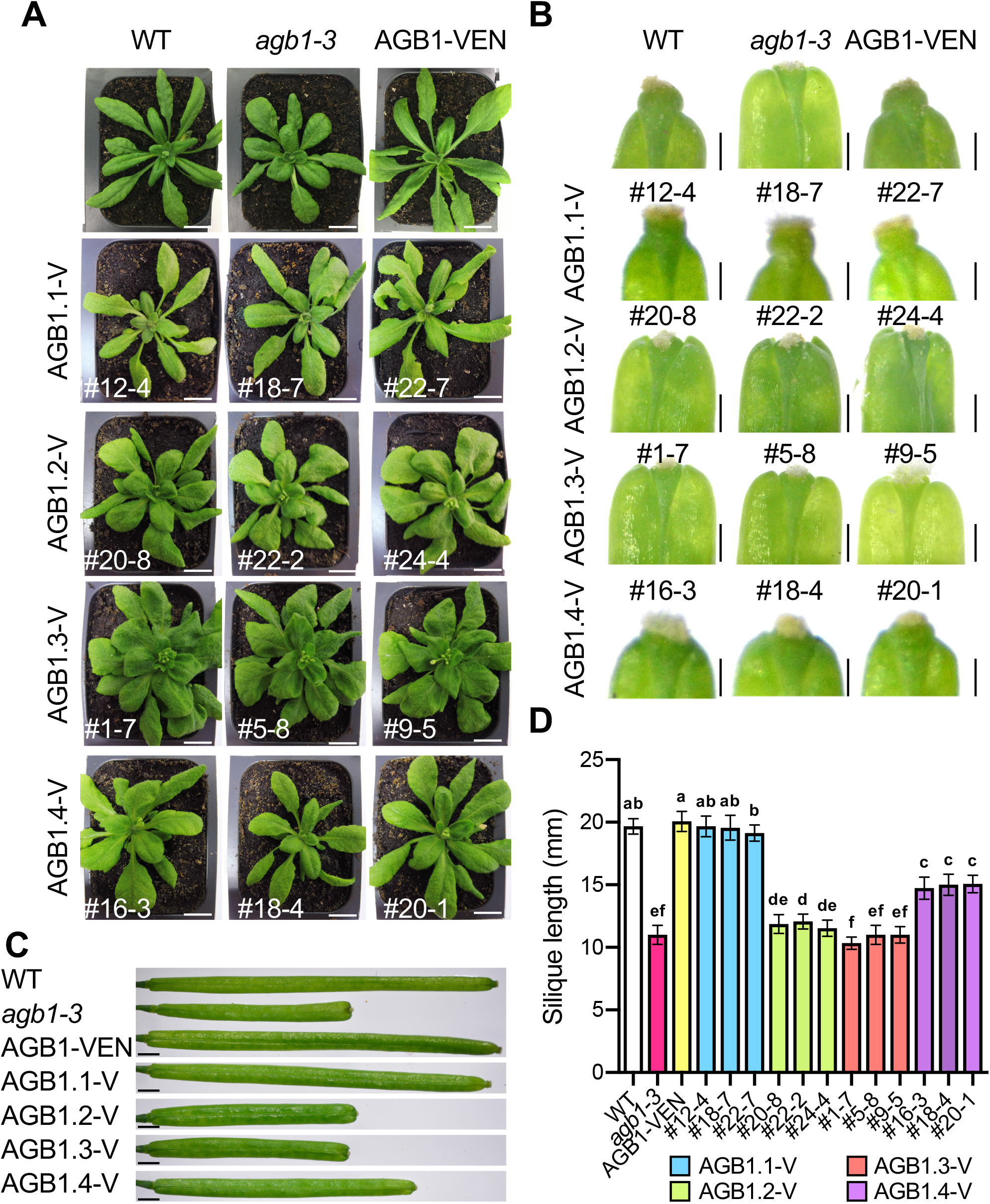
Morphological observations of 4 AGB1 variants. Morphological observation of wild-type (WT, white), *agb1-3* (pink), ProAGB1:AGB1-Ven a*gb1-3* line #6-3 (AGB1-VEN, yellow), ProAGB1:AGB1.1-Ven *agb1-3* line #12-4, #18-7, #22-7 (AGB1.1-V, sky blue), ProAGB1:AGB1.2-Ven *agb1-3* line #20-8, #22-2, #24-4 (AGB1.2-V, light green), ProAGB1:AGB1.3-Ven *agb1-3* line #1-7, #5-8, #9-5 (AGB1.3-V, orange red), ProAGB1:AGB1.4-Ven *agb1-3* line #16-3, #18-4, #20-1 (AGB1.4-V, light purple) grown on soil for 24 days (A) or 42 days (B, C, D). Representative images are shown for rosette leaves (A), silique tip (B) and mature silique (C) from at least 12 plant per lines. (D) Quantitative analysis of silique length obtained from 16 siliques shown in mean ± SD (mm). Different letters indicate significant difference at P < 0.05 as determined by one-way ANOVA. Values with the same letter are not significantly different. Scale bars, 1 cm (A) and 3 mm (B, C).

### The splice isoforms of AGB1 differentially complement defects in ER stress

To test which AGB1 splice isoform is required for ER stress tolerance, we performed TM-sensitivity assay by using AGB1.1-V (lines #12-4, #18-7, #22-7), AGB1.2-V (lines #20-8, #22-2, #24-4), AGB1.3-V (lines #1-7, #5-8, #9-5) and AGB1.4-V (lines #16-3, #18-4, #20-1) with WT, *agb1-3* and AGB1-VEN. For seedlings grown on MS-agar plates for 2 weeks (mock), the fresh weight was greater for *agb1-3*, AGB1.2-V and AGB1.3-V than the WT (Figure 6A and B). Under ER stress (TM), *agb1-3* exhibited severe growth defects as compared with the WT, whereas AGB1.1-V fully rescued this phenotype, which suggests that AGB1.1-V may be necessary and sufficient for ER stress tolerance. In addition to AGB1.1-V, AGB1.4-V showed partial complementation, but AGB1.2-V and AGB1.3-V failed to rescue it (Figure 6). Likewise, *AGB1.1* was necessary and sufficient to rescue phenotypes in development and *AGB1.4* could partially complement them, whereas *AGB1.2* and *AGB1.3* failed in ER stress tolerance (Figure 6A, 6B).

**Fig. 6.**
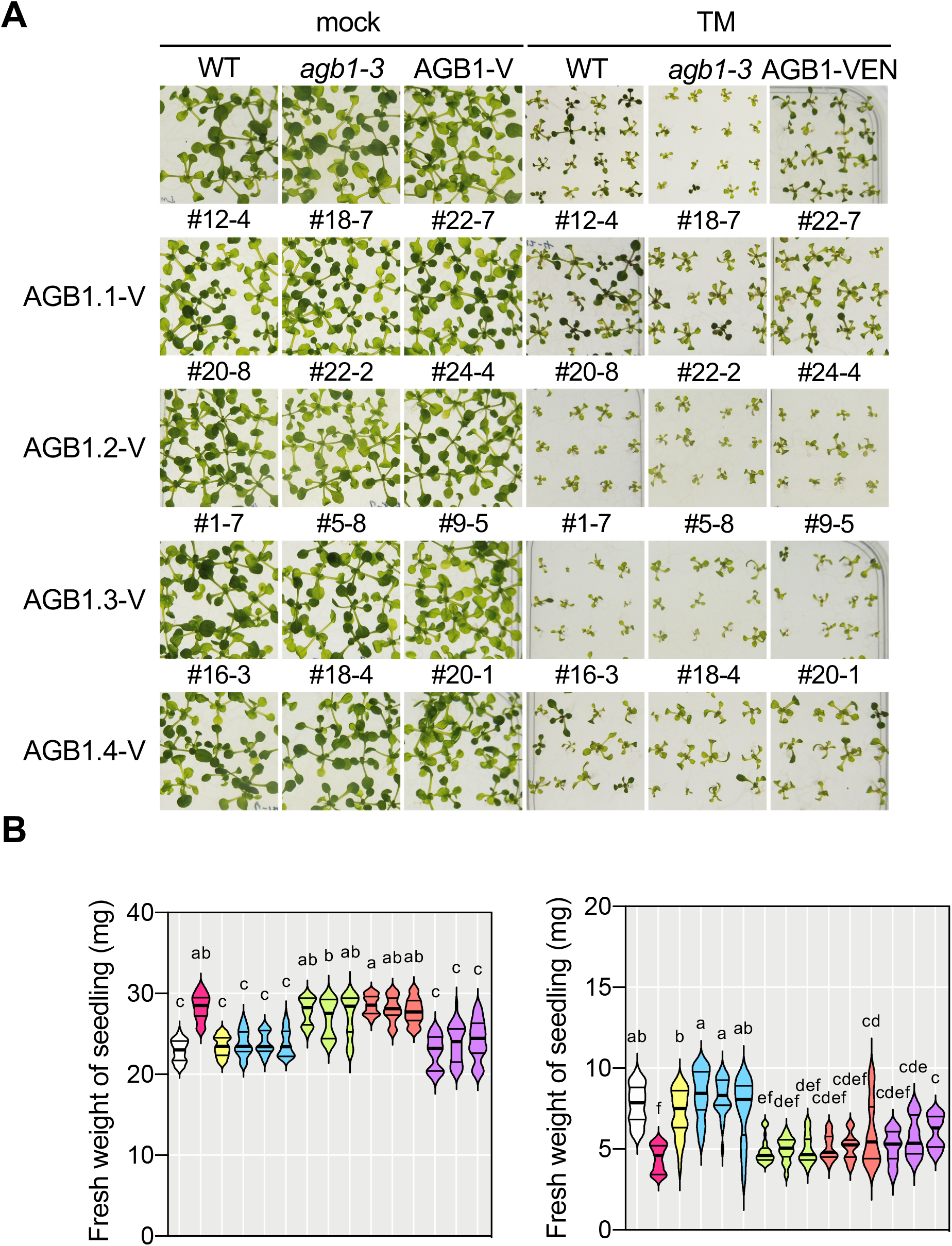
Functional complementation of hypersensitivity to ER stress in *agb1-3* mutant by 4 AGB1 variants. Representative images (A) and fresh weight (B) of 14-day-old of wild-type (WT, white), *agb1-3* (pink), ProAGB1:AGB1-Ven *agb1-3* line #6-3 (AGB1-VEN, yellow), ProAGB1:AGB1.1-Ven *agb1-3* line #12-4, #18-7, #22-7 (AGB1.1-V, sky blue), ProAGB1:AGB1.2-Ven *agb1-3* line #20-8, #22-2, #24-4 (AGB1.2-V, light green), ProAGB1:AGB1.3-Ven *agb1-3* line #1-7, #5-8, #9-5 (AGB1.3-V, orange red), and ProAGB1:AGB1.4-Ven *agb1-3* line #16-3, #18-4, #20-1 (AGB1.4-V, light purple) plants grown on 1/2 MS containing dimethyl sulfoxide (mock in A, left panel in B) or tunicamycin (TM in A, right panel in B) are shown. Fresh weight of the seedlings in A was measured individually (n=16), and shown as mean ± SD (mg) in B. Different letters indicate significant difference at P < 0.05 as determined by one-way ANOVA. Values with the same letter are not significantly different. Three biologically independent experiments were performed with similar results.

### AGB1 splicing isoforms are localized differentially in cells

AGB1.1-V and AGB1.4-V complemented the defects in both plant development and ER stress tolerance (Figures 5 and 6), whereas AGB1.3-V enhanced the phenotype of rosette leaves (Figure 5A). To test whether a difference in subcellular localization causes distinct functions *in vivo*, we observed fluorescent signals in root epidermis of stable transgenic plants (Figure 7A). For AGB1.1-V, the Venus signal was merged with ER-tracker and FM4-64 (Figure 7A), which indicates that AGB1.1-V is localized at the PM and ER similar to AGB1-VEN (Figure 2). In contrast, the signals of AGB1.3-V and AGB1.4-V were barely detected (Figure 7A) even though the expression was higher for *AGB1.4* than *AGB1.1* (Figure 4). To further investigate the subcellular localization, we used a protoplast transient assay with the AGB1 isoforms fused C-terminally to mRFP expressed by the cauliflower mosaic virus 35S promoter (Pro35S: AGB1.x-mRFP, Figure 7B, 7C and 7D) together with organellar markers. AGB1.1-mRFP and AGB1.4-mRFP overlapped with both the PM marker (PM-yk in Figure 7B) and ER marker (ER-gk, Figure 7C), so both isoforms were located at the PM and ER. Unlike AGB1.1-mRFP and AGB1.4-mRFP, AGB1.3-mRFP showed a dot-like structure that overlapped with neither PM-yk nor ER-gk (Figure 7B and C). To test whether AGB1.3-V is localized at the nucleus, we stained protoplasts expressing AGB1.3-mRFP with DAPI. Because both signals partially overlapped, AGB1.3-V might be localized at the nucleus (Figure 7D). In conclusion, AGB1.1 and AGB1.4 may be localized mainly at the PM and ER, whereas AGB1.3 may be at the nucleus.

**Fig. 7.**
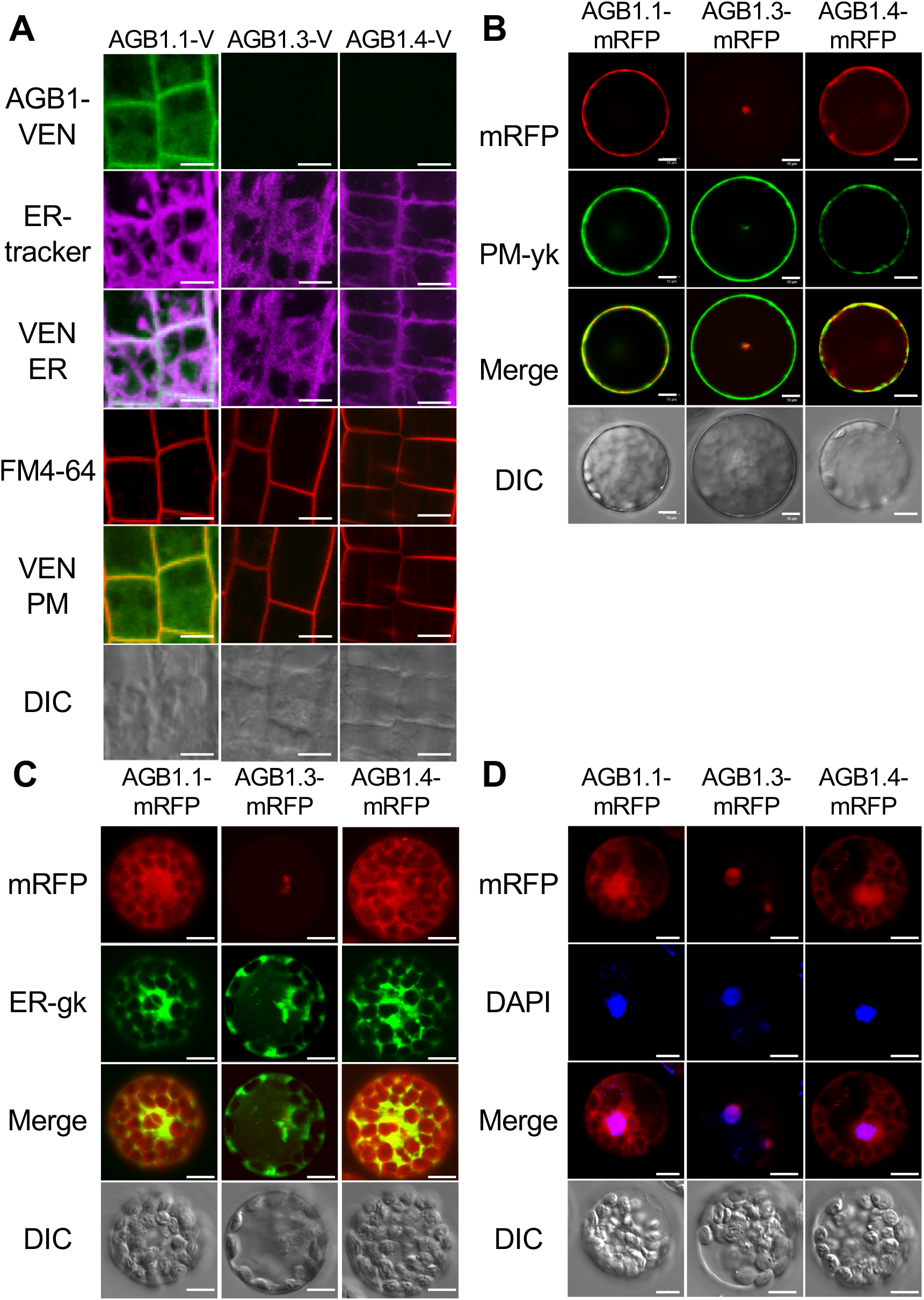
The subcellular localization of AGB1.x-V. Representative images of AGB1.x-V in ProAGB1:AGB1.x-Ven *agb1-3* plants (AGB1.1-V, #12-4; AGB1.3-V, #5-8; AGB1.4-V, #18-4) (A) and in wild-type protoplasts transiently expressing ProAGB1:AGB1.x-Ven (B, C, D). (A) Subcellular localization of AGB1.x-V in root of 7-day-old AGB1.x-V plants. Fluorescence of AGB1.x-V (VEN, Green) and co-staining of the ER by ER-tracker (ER, Magenta) and the plasma membrane by FM4-64 (PM, red). Cell structure is shown as differential interference contrast (DIC) image at the bottom. (B, C) Pro35S:AGB1.x-mRFP (mRFP, red) was co-transduced with the plasma membrane marker (PM, green, PM-yk in B) or the ER marker (ER, green, ER-gk in C) into wild-type protoplast. (D) wild-type protoplast expressing Pro35S:AGB1.x-mRFP (mRFP, red) was staining with DAPI (nucleus, blue, in D). Co-localization signals in (A) of AGB1.x-V and ER-tracker or plasma membrane staining dye FM4-64 or (B, C, D) AGB1-mRFP with marker of ER, plasma membrane and nuclear staining dye are shown in merged images (A: VEN/ER and VEN/PM; B: mRFP/PM-yk; C: mRFP/ER-gk; D: mRFP/DAPI). Scale bars, 10 µm.

### AGB1.3 does not interact with AGGs

To investigate how an interaction with Gγ components affects functions of AGB1 splice isoforms, we used a yeast two-hybrid assay with full-length coding sequences of AGB1.1, AGB1.3 and AGB1.4 as bait and 3 AGGs, AGG1, AGG2 and AGG3, as prey. We cultured yeast diploid cells, then serially diluted them before spotting cells on a synthetic double dropout plate (SD-Leucine-Tryptophan, DDO). All cells including the positive (p53/T) and negative control cells (Lam/T) equally grew on DDO plates, so that all yeast strains expressed both bait and prey. To test an interaction of each combination, we grew cells on DDO plates containing X-α-gal and aureobasidin A (DDOX/A). AGB1.1 and AGB1.4 showed interactions with all AGGs, with no interactions found for AGB1.3 (Figure 8). These results were confirmed by an additional selection plate, triple dropout (TDO, DDO-Histidine). Notably, the interaction between AGG2 and both AGB1.1 and AGB1.4 was weaker than that with AGG1 and AGG3. In conclusion, both AGB1.1 and AGB1.4 interacted with AGG1, AGG2 and AGG3, but AGB1.3 did not interact with any AGG components in the yeast two-hybrid assay.

**Fig. 8.**
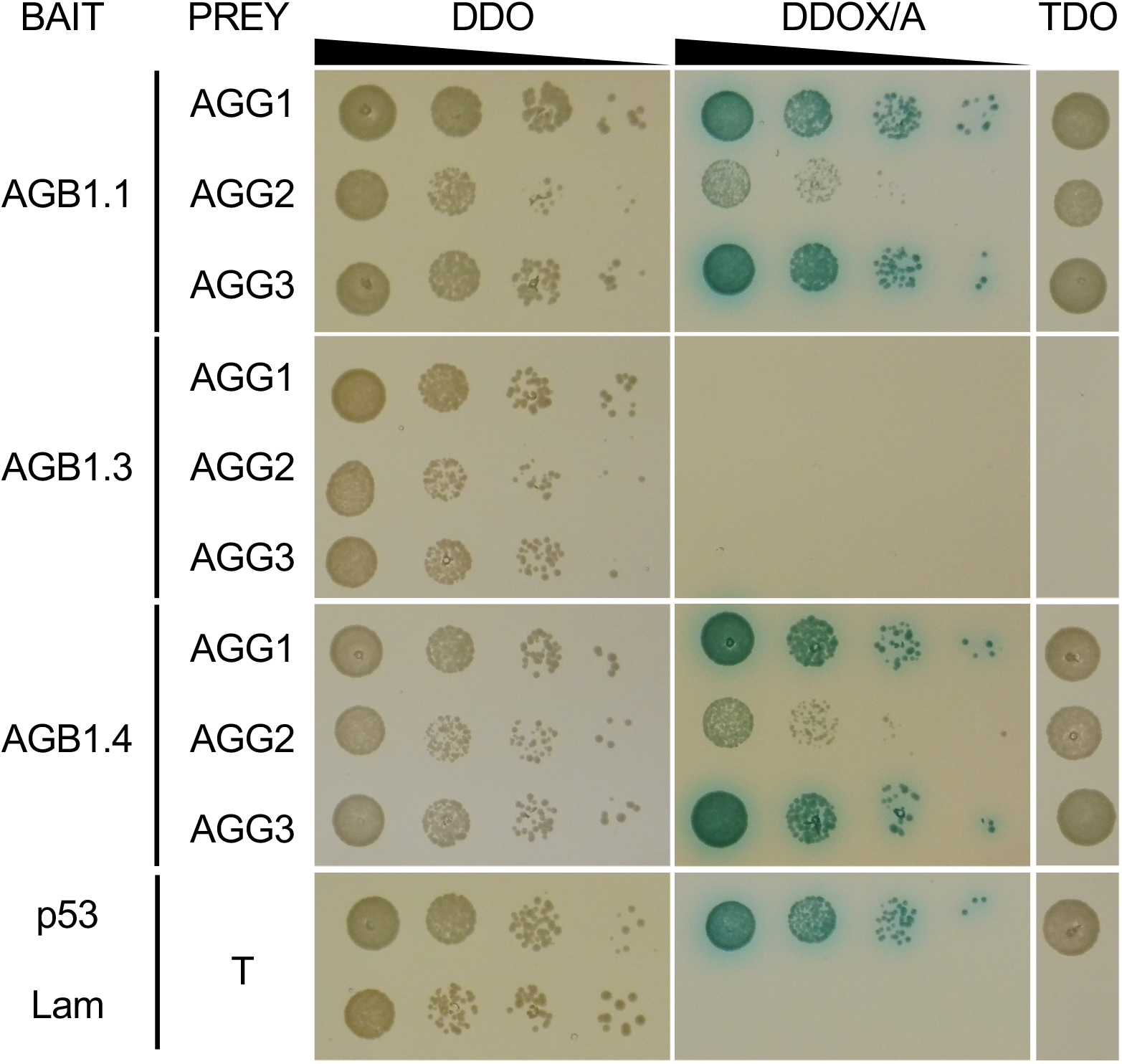
Protein-protein interactions of AGB1 splicing variants with AGG proteins in yeast. Yeast 2-hybrid (Y2H) interactions between AGB1 variants and AGGs. The culture of yeast diploid cells bearing the vectors indicated were serially diluted (10-fold dilution from left to right) and spotted on control media (DDO, SD-Leucine-Tryptophan), media containing X-α-gal and Aureobasidin (DDOX/A), and media lacking histidine (TDO, SD-Leu-Trp-His) on agar plates. The combination of p53-BD and T-AD is shown as a positive control, with Lam-BD and T-AD combination as a negative control. The cells were incubated for 2 days at 30 °C.

## DISCUSSION

The functions of the heterotrimeric G protein complex in plants have been extensively studied for a few decades because early genetic studies suggested its pivotal roles in plant development. In Arabidopsis, knock-out mutations of *AGB1* cause diverse phenotypes in both plant development and stress responses. Although previous studies using different methods suggested multiple subcellular localization of AGB1 protein at the PM, ER, and nucleus, no conclusive evidence has been provided regarding the functional relevance of a differential subcellular localization of AGB1 proteins in a whole plant body. To reveal the precise molecular mechanisms of how the sole component of the Gβ subunit, AGB1, plays diverse roles *in vivo*, we noted that the splice variants of *AGB1* produce 4 isoforms of AGB1 protein with distinct subcellular localization.

Here, we successfully established stable transgenic lines expressing AGB1-VEN under its own promoter (*ProAGB1:AGB1-Ven*, *agb1-3*), which was able to rescue the phenotypes in both plant development and stress tolerance (Figure 1). AGB1-VEN was localized at the PM and ER (Figure 2). The ER was an unexpected location because AGB1 was predicted to be at the PM by binding with Gγ subunit like animal cells. The early study using Arabidopsis wild-type plants expressing the cDNA of AGB1 (corresponding to AGB1.1) fused to GFP driven by the 35S promoter showed AGB1-GFP localized to the PM and the nucleus in leaf epidermal cells of stable transgenic lines (Anderson and Botella, 2007). This discrepancy may be due to the different promoters used: 35S in (Anderson and Botella, 2007) and its own promoter in the current study. In fact, the other study using the *ProAGB1:CFP-AGB1* construct supported the ER localization of AGB1 by membrane fractionation (Wang et al., 2007). AGB1 proteins driven by their own promoter are most likely localized at the PM and ER. The ER localization seems reasonable because AGB1 plays a role in ER stress (Chen and Brandizzi, 2012, Cho et al., 2015). In our transgenic plant AGB1-VEN, one splice isoform, AGB1.3, was expressed without the VEN reporter because of the termination of translation in the 7th WD40 motif (Figure 3B, 3C). This observation may account for why the VEN signal in AGB1-VEN plants shown in Figure 2 represents the subcellular localization of AGB1.1, AGB1.2, and AGB1.4 but not AGB1.3.

Our comprehensive study of the 4 splice isoforms of AGB1 revealed that *AGB1.1* was necessary and sufficient to rescue the *agb1-3* phenotypes. *AGB1.1* produced the full-length protein localized at the PM and ER in root (Figure 7A). Previous studies suggested the function of AGB1 in the nucleus (Xu et al., 2015, Xu et al., 2017, Zhang et al., 2017, Lian et al., 2018, Xu et al., 2019), so nuclear localization of AGB1.1-VEN cannot be ruled out despite our failure to observe this in our experimental conditions (Figure 7A). Although the transcript level was higher for *AGB1.4* than *AGB1.1* in all wild-type tissues (Figure 4), AGB1.4-V was undetectable in roots of 7-day-old seedlings (Figure 7A), which suggests a possible posttranscriptional regulation of *AGB1.1* and *AGB1.4*. AGB1.4-V protein might be unstable owing to the loss of 5 amino acids, VARYS, in the coiled-coil region as compared with the full-length AGB1.1-V. Indeed, AGB1.4 complemented the *agb1-3* phenotypes partially but not fully in both development and ER stress tolerance (Figures 5D and 6B). The protein level of AGB1.4-V may not be high enough to fully complement the *agb1-3* phenotypes. The remaining 2 splice isoforms, AGB1.2 and AGB1.3, were not functional because both failed to rescue the *agb1-3* phenotypes (Figures 5 and 6). We cannot exclude that these two proteins are not functional in vivo because the VEN reporter protein simply inhibits the functions of AGB1.2 and AGB1.3. However, the complementation assay suggested that AGB1.3-VEN enhanced the leaf phenotype of *agb1-3* (Figure 5A), so AGB1.3 may show a dominant negative effect during leaf development. Furthermore, unlike AGB1.1 and AGB1.4, AGB1.3 may be localized in the nucleus (Figure 7D). Whether AGB1.3 plays a role in the nucleus to show a dominant negative effect awaits future investigation.

The yeast 2-hybrid assay revealed that AGB1.1 and AGB1.4 physically interact with AGG1, AGG2, and AGG3, with AGG2 interacting weaker than AGG1 or AGG3 (Figure 8). The interaction between AGB1 with AGGs has been shown by a couple of studies, but our results newly revealed the following 4 points: 1) the interacting ability of AGB1 with AGGs is distinct among splicing isoforms of AGB1; 2) as compared with AGG1 and AGG3, AGG2 weakly interacts with AGB1.1 and AGB1.4; 3) because of no differences between AGB1.1 and AGB1.4, a loss of 5 amino acid residues in the coiled-coil domain in AGB1.4 has no effect with respect to the interaction of AGGs; and 4) the 7th-WD40 motif is necessary for interaction between AGB1 and AGGs because AGB1.3 failed to interact with all AGG isoforms (Figure 8).

In summary, the current study revealed functional relevance of 4 splice isoforms of AGB1 at the protein level. The transgenic lines established and the resulting functional data help decode the complex molecular mechanism underlying the multifaceted function of Arabidopsis Gβ protein in plant growth regulation and stress response.

## MATERIAL AND METHODS

### Plant materials and growth conditions

*Arabidopsis thaliana* plants were in Columbia-0 (Col-0) ecotype and were grown under a 16-h light/8-h dark photoperiodic condition at 22 °C with light intensity 150 µmol min^-1^ sec^-1^. Arabidopsis seeds were surface-sterilized with 70% ethanol for 20 min, then rinsed with sterile distilled water 5 times. Surface-sterilized seeds were kept at 4 °C for 1 day to synchronize germination. Cold-stratified seeds were sown onto half-strength Murashige and Skoog (1/2 MS) medium (pH 5.6) with 0.6% (w/v) agar and 0.86% (w/v) sucrose (Murashige and Skoog, 1962) or in a soil pot. The isolation of the *agb1-3* mutant (SALK_061896) was described previously (Cho et al., 2015).

### Phenotype quantification

For plate-based analysis, the imbibed seeds were observed upon transfer to a growth chamber for the time indicated. For soil-based analysis, stratified seed were observed upon transfer to growth chamber for the time indicated. The rosette leave morphology was compared between lines before bolting at 18 days old. The silique morphology was compared for the longest siliques from the primary inflorescence stem from 42-day-old plants. All phenotypic observations were performed with at least 12 plants for each line per biological replicate. The quantitative data were obtained from 3 biologically independent experiments.

### Complementation assay of hypersensitivity to tunicamycin (TM)-induced ER stress of *agb1-3*

Plants were grown on 1/2 MS agar plates containing 0.01% (v/v) dimethyl sulfoxide (mock) or 50 ng mL-1 TM (654380; Merck, Darmstadt, Germany) for 14 days, and photographed. Fresh weight of 14-day-old seedlings was measured individually (n=25 or 16), and shown as mean ± SD (mg). Three biologically independent experiments were performed.

### Vector construction and Arabidopsis transgenic lines

#### *ProAGB1: AGB1-Ven agb1-3* (AGB1-VEN transgenic lines)

The *Sfo*I site was added immediately before the stop codon for AGB1 of pCC83 (*pENTR-ProAGB1:AGB1*) (Cho et al., 2015) by PCR-based site-directed mutagenesis (Sawano and Miyawaki, 2000) with the primer KK235 to obtain pCC87 (*pENTR-ProAGB1:AGB1-Sfo*I) for creating the fluorescent reporter Venus (Ven) fused translationally to the C-terminus of the open reading frame of AGB1. The obtained pCC91 (*pENTR-ProAGB1:AGB1-Ven*) was then recombined into a pBGW destination vector (Karimi et al., 2002) by using Gateway LR Clonase II (Invitrogen, Thermo Fisher Scientific, Waltham, MA, USA) to obtain pCC95 (*pBGW-ProAGB1:AGB1-Ven*). pCC95 was transduced into *agb1-3* via *Agrobacterium tumefaciens* strain GV3101-mediated transformation. In total, 24 T1 plants were selected on soil by spraying with 0.1% BASTA solution. The obtained T2 seeds were screened by using BASTA and selected by genotyping. To distinguish transgenic *ProAGB1:AGB1-Ven* from endogenous *AGB1*, specific primers (KK212/KK104) were designed. Lines #6-3, #10-1, #17-2 were used for observation and line #6-3 was a representative line.

#### *ProAGB1: AGB1.x-Ven agb1-3* (AGB1.x-V transgenic lines)

The genomic fragment from the transcription start site of AGB1 to the *Sfo*I site immediately before the Venus fragment was deleted from pCC91 (*pENTR-ProAGB1:AGB1-Ven*) by PCR-based site-directed mutagenesis (Sawano and Miyawaki, 2000) with the primer KK638 containing *BamH*I site to obtain pYC26 (*pENTR-ProAGB1:Ven*). The coding sequences of *AGB1.1*, *AGB1.2* and *AGB1.3* variants were amplified and generated with the *BamH*I sites before the start codon and stop codon with the primer set YC12/YC14, YC13/YC14 and YC12/YC15 to obtain pYC31 (*pENTR-ProAGB1:AGB1.1-Ven*), pYC32 (*pENTR-ProAGB1:AGB1.2-Ven*) and pYC37 (*pENTR-ProAGB1:AGB1.3-Ven*). For *AGB1.4*, 15 nucleotides were deleted from pYC31 by PCR-based site-directed mutagenesis (Sawano and Miyawaki, 2000) with the primer YC24 to obtain pYC35 (*pENTR-ProAGB1:AGB1.4-Ven*). The obtained entry vector plasmids pYC31, pYC32, pY37 and pYC35 were recombined into the pBGW destination vector (Karimi et al., 2002) by use of Gateway LR reaction to obtain pYC33 (*pBGW-ProAGB1:AGB1.1-Ven*), pYC34 (*pBGW-ProAGB1:AGB1.2-Ven*), pYC38 (*pBGW-ProAGB1:AGB1.3-Ven*), pYC36 (*pBGW-ProAGB1:AGB1.4-Ven*), respectively. pYC33, pYC34, pYC38, and pYC36 were transduced into *agb1-3* via *Agrobacterium tumefaciens* strain GV3101-mediated transformation. In total, 48 T1 plants were selected on soil by spraying with 0.1% BASTA solution. The obtained T2 seeds were screened by using BASTA and selected by genotyping. To distinguish transgenic *ProAGB1:AGB1.x-Ven* from endogenous *AGB1*, specific primers (KK212/KK104) were designed for genotyping. Lines #12-4, #18-7, #22-7 were used for observing AGB1.1-V, and #12-4 was a representative line. Lines #20-8, #22-2, #24-4 were used for observing AGB1.2-V, and #24-4 was a representative line. Lines #1-7, #5-8, #9-5 were used for observing AGB1.3-V, and line #5-8 was a representative line. Lines #16-3, #18-4, #20-1 were used for observing AGB1.4-V, and line #18-4 was representative line.

#### Pro35S:AGB1.x-mRFP

The protein coding sequences of *AGB1.1* and *AGB1.3* were amplified by PCR with the primer sets YC290/YC490 and YC493/YC494, respectively. The fragments were cloned into the pENTR/D-TOPO plasmid vector (Invitrogen) to obtain pYC19 (*pENTR-AGB1.1*) and pYC28 (*pENTR-AGB1.3*), respectively. For *AGB1.4*, 15 nucleotides were deleted from pYC19 by PCR-based site-directed mutagenesis (Sawano and Miyawaki, 2000) with the primer YC24 to obtain pYC30 (*pENTR-AGB1.4*). The obtained entry vector plasmids pYC19, pYC28 and pYC30 were recombined into the pGWB654 destination vector (*pPZP-Pro35S:C-mRFP*) (Karimi et al., 2002) by using Gateway LR reaction to obtain pYC84 (*Pro35S:AGB1.1-mRFP*), pYC90 (*Pro35S:AGB1.3-mRFP*) and pYC91(*Pro35S:AGB1.4-mRFP*), respectively.

#### mRFP-HDEL

To generate the fluorescent ER marker, the gene encoding mRFP (675 bp) was first amplified by using the pGWB653 vector (pPZP_C-mRFP) with oligonucleotides (YC331/YC332), then PCR products (half-length of pumpkin 2S albumin signal peptide-mRFP-ER retention signal) were used as a template with oligonucleotides (YC334/YC332) to obtain a chimeric gene encoding SP-mRFP-HDEL (786 bp, pumpkin 2S albumin signal peptide-mRFP-ER retention signal) by PCR mutagenesis (Sawano and Miyawaki, 2000). The SP-mRFP-HDEL fragments were cloned into the pENTR/D-TOPO vector as pYC62 (*pENTR:SP-mRFP-HDEL*) and recombined into a pGWB602 destination vector (pPZP_35S pro) (Karimi et al., 2002) by using Gateway LR Clonase II (Invitrogen) to obtain pYC65 (*Pro35S:SP-mRFP-HDEL*).

#### pGBKT7-AGB1.x *and* pGADT7-AGGs

To clone *AGB1.1*, *AGB1.3* and *AGB1.4*, their protein coding regions (1,134, 1033 and 1,119 bp, respectively) were amplified by PCR from vector plasmids pYC19, pYC28 and pYC30 by using Phusion high-fidelity DNA polymerase (F530S, Invitrogen) with the primer sets YC269/YC270, YC269/YC582 and YC269/YC270 to create restriction enzyme sites, respectively. For cloning *AGG1*, *AGG2* and *AGG3*, protein coding regions (297, 303 and 756 bp, respectively) were amplified by PCR from Arabidopsis wild-type seedling cDNA, by using Phusion high-fidelity DNA polymerase with the primer sets YC339/YC342, YC585/YC586 and YC587/YC588, respectively. PCR products were digested with restriction enzymes *Eco*RI and *Xho*I for AGG1, *Nde*I and *BamH*I for *AGB1.1*, *AGB1.3*, *AGB1.4*, *AGG2* and *AGG3*. The PCR fragments of AGB1.x were ligated into the pGBKT7 DNA BD vectors (Takara Bio USA, CA, USA) and AGGs into the pGADT7 AD vector (Takara Bio) to produce pYC59 (*pGBKT7-AGB1.1*), pYC95 (*pGBKT7-AGB1.3*), pYC98 (*pGBKT7-AGB1.4*), pYC58 (*pGADT7-AGG1*), pYC96 (*pGADT7-AGG2*) and pYC97 (*pGADT7-AGG3*) for expression in *Saccharomyces cerevisiae* strain Y2HGold or Y187 (Takara Bio).

### Confocal microscopy observation for in planta fluorescent imaging

For imaging in primary root, Venus fluorescence in 7-day-old seedlings of AGB1-VEN, AGB1.1-V, AGB1.3-V and AGB1.4-V were observed at the root epidermis under a confocal laser-scanning microscope (LSM 510 Meta; Carl Zeiss, Jena, Germany). To observe subcellular localization, seedling samples were immersed in 10 µg/ml FM4-64 (T13320, Invitrogen) for 3 min to stain plasma membrane, 1 µM ER-tracker Blue-White DPX (E12353, Invitrogen) for 5 min to stain endoplasmic reticulum.

For transient expression in Arabidopsis protoplasts, a drop of transformed protoplasts was applied onto a glass slide with a ring sticker. The fluorescent signals were observed under a confocal laser-scanning microscope (LSM 510 Meta). To observe subcellular localization, the plasmid pYC84 (*Pro35S:AGB1.1-mRFP*), pYC90 (*Pro35S:AGB1.3-mRFP*) or pYC91 (*Pro35S:AGB1.4-mRFP*) were co-transformed with the plasma membrane marker PM-yk, endoplasmic reticulum ER-gk (Nelson et al., 2007) or stained with 2 µg mL^-1^ DAPI (D1306; Invitrogen) to stain nuclei. After staining, the samples were observed under a confocal microscope and images were captured by using LSM 510 v3.2 (Carl Zeiss) with filters for DAPI or ER-tracker Blue-White DPX (Diode 405 nm laser, band-pass 420-480 nm); ER-gk (Argon 488 nm laser, band-pass 505-530 nm); Venus, PM-yk (Argon 514 nm laser, band-pass 520-555 nm); mRFP (HeNe 543 nm laser, band-pass 560-615 nm); or FM4-64 (HeNe 543 nm laser, long-pass 650 nm).

### Transient expression assay

Arabidopsis protoplast isolation and transformation were conducted as reported in Wu et al. (Wu et al., 2009).

### Yeast strains and yeast 2-hybrid assay

The pGBKT7 plasmids were transduced into S. cerevisiae Y2HGold strain to produce the strains YCY10 (Y2HGold/pGBKT7), YCY12 (Y2HGold/pYC59, AGB1.1-BD), YCY13 (Y2HGold/pYC96, AGB1.2-BD), YCY14 (Y2HGold/pYC97, AGB1.3-BD), and YCY15 (Y2HGold/pYC98, AGB1.4-BD). The pGADT7 plasmids were transformed into *S. cerevisiae* Y187 strain to produce the strains YCY11 (Y187/pGADT7), YCY16 (Y187/pYC58, AGG1-AD), YCY17 (Y187/pYC94, AGG2-AD), and YCY18 (Y187/pYC95, AGG3-AD). The plasmid transduction into the above-mentioned yeast strains was confirmed by PCR (KK70/CY179 for AGB1.x-BD, KK281/CY180 for AGG1-AD, YC265/CY180 for AGG2-AD, KK297/CY180 for AGG3-AD), and the oligonucleotide sequences are listed in Supplemental Table S1. The yeast diploid cells were generated by crossing yeast strain Y2HGold with Y187 as indicated and then sprayed on a synthetic dropout-Leucine-Tryptophan selection plate (DDO). For yeast two-hybrid assay, the yeast diploid cells were grown logarithmically at 30 °C in DDO media; 5 µl of OD_600_ = 0.1 culture was spotted and underwent 10-fold serial dilutions on a DDO plate, synthetic complete-Leucine-Tryptophan-Histidine selection plate (TDO), and synthetic-Leucine-Tryptophan with 5-Bromo-4-Chloro-3-Indolyl-α-D-Galactopyranoside (X-α-gal) and Aureobasidin A (Aba) selection plate (DDOX/A) and were incubated for 48 h at the 30 °C. For control, the diploid cells harbored pGBKT7-p53 and pGADT7-T (YCY19) as a positive set and the diploid cells harbored pGBKT7-Lam and pGADT7-T (YCY20) as a negative set.

### RNA preparation, reverse transcription (RT) and RT-qPCR

Seedlings were frozen by immersion in liquid nitrogen and stored at -80 °C until use. Total RNA from 10 of 7-day-old seedlings was extracted by using TRI reagent (AM9738, Invitrogen). In total, 500 ng RNA was used for complementary DNA (cDNA) synthesis by using the SuperScript III First-Strand Synthesis SuperMix (11752050, Invitrogen). In total, 50 ng cDNA was used as a template for RT-qPCR using the 7500 fast real-time PCR system (Applied Biosystems). Data were analyzed by the comparative threshold cycle method (ΔCT methods). The transcript level was normalized to that of *ACTIN2* gene (*ACT2*, KK129/KK130) for each sample. For *AGB1.1* (YC89/YC90), *AGB1.2* (YC91/YC92), *AGB1.3* (YC93/YC94), *AGB1.4* (YC95/YC99), the transcript level was shown as the 2^ΔCT^ (Mean + SD) from 3 biological replicates with 3 technical replicates. The primer sets for RT-qPCR are shown in Supplemental Table S1.

### Statistical analysis

Statistical analyses were performed with RSTUDIO 1.3.1093 (Rstudio Team, 2020). Data obtained from at least 3 biologically independent experiments were analyzed by one-way ANOVA. Statistically different groups among conditions were further evaluated for significance with the Tukey’s honestly significant difference post-hoc test and displayed with different letters indicating means that differ significantly. P < 0.05 was considered statistically significant.

### Accession Numbers

Sequence data from this article can be found in the GenBank/EMBL data libraries under accession numbers At4g34460 (*AGB1*), At3g63420 (*AGG1*), At3g22942 (*AGG2*), and At5g20635 (*AGG3*).

## ACKNOWLEDGEMENTS

The author thanks Chia-En Chen (Institute of Plant and Microbial Biology, Academia Sinica, Taiwan) for technical assistance with molecular cloning and Kazue Kanehara (Institute of Plant and Microbial Biology, Academia Sinica, Taiwan) for comment on the article.

## AUTHOR CONTRIBUTIONS

Y. C. designed research, performed experiments, analyzed data and wrote the manuscripts.

## SUPPORTING INFORMATION

**Fig. S1.**
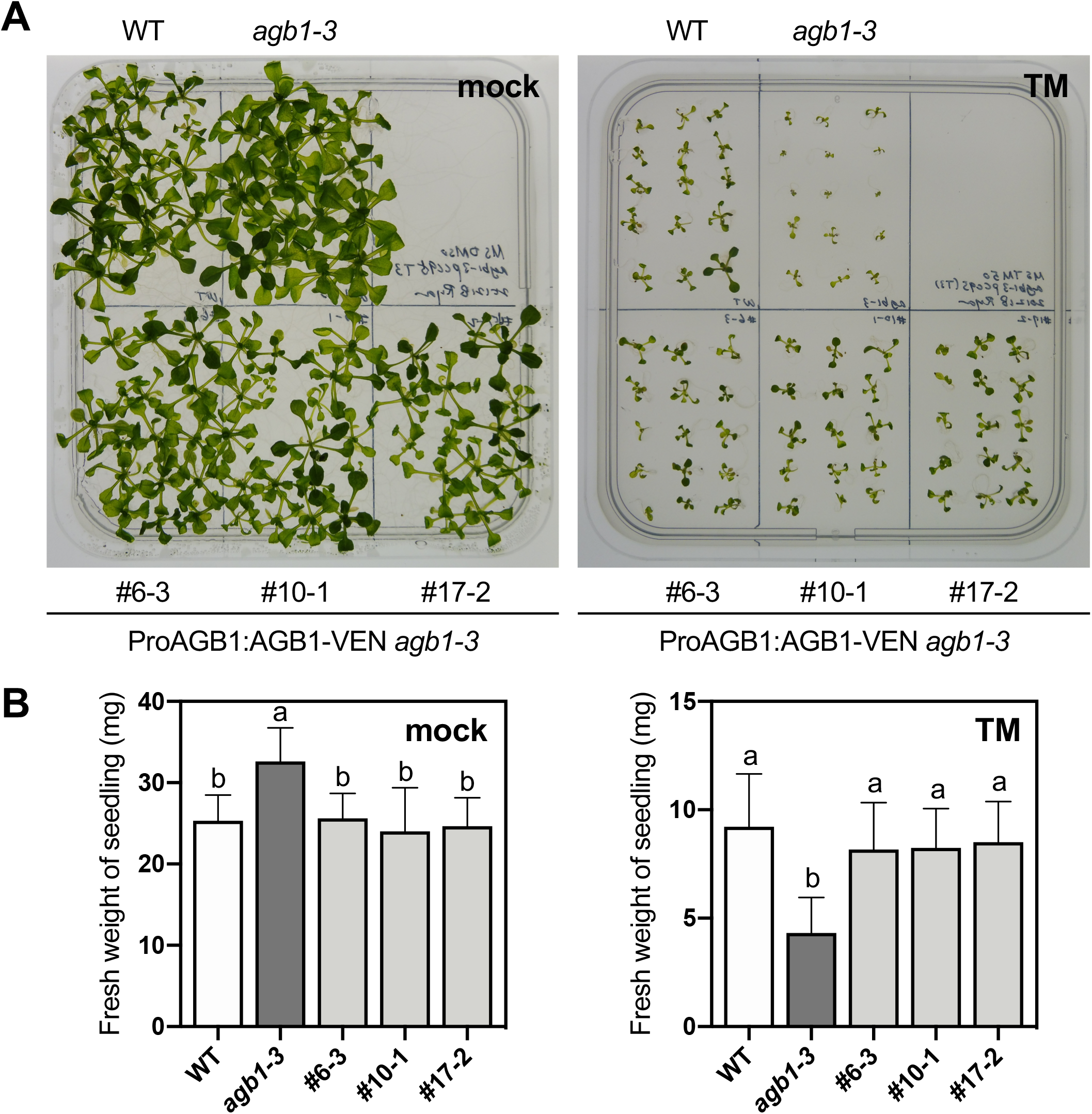
Functional complementation of hypersensitivity to ER stress in *agb1-3* mutant by the other lines of AGB1-VEN transgenic plants. Representative images (A) and fresh weight (B) of 14-day-old wild-type (WT), *agb1-3* mutant and AGB1-VEN (ProAGB1:AGB1-Ven *agb1-3*; #6-3, #10-1, #17-2) plants grown on 1/2 MS containing dimethyl sulfoxide (mock) or tunicamycin (TM) are shown. Fresh weight of the seedlings in (A) was measured individually (n=25), and shown in mean ± SD (mg) in (B). Different letters indicate significant difference at P < 0.05 as determined by one-way ANOVA. Values with the same letter are not significantly different. Three biologically independent experiments were performed with the similar results.

**Fig. S2.**
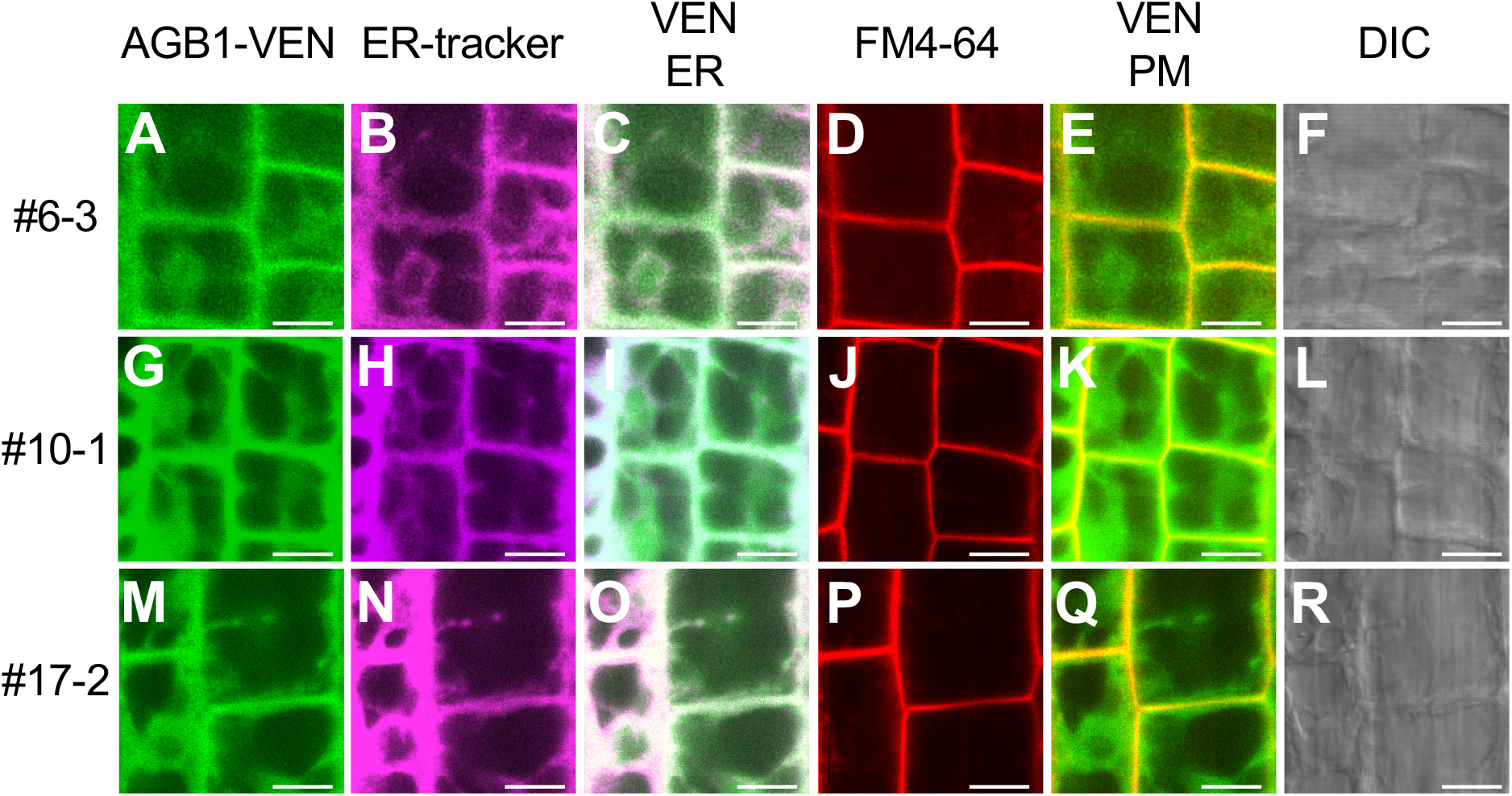
Subcellular localization of AGB1 in the other lines of AGB1-VEN transgenic plants. Representative images of ProAGB1:AGB1-Ven (AGB1-VEN) in roots of 7-day-old ProAGB1:AGB1-Ven *agb1-3* plants (lines #6-3, #10-1, #17-2) are shown. Fluorescence of AGB1-VEN (Green, A, G, M) and co-staining of the ER by ER-tracker (ER; magenta, B, H, N), plasma membrane by FM4-64 (PM; Red, D, J, P) are shown. Co-localization signals of AGB1-VEN and ER or plasma membrane staining dye are shown in merged images (C, I, O) or (E, K, Q), respectively. Differential interference contrast (DIC) images are shown on the right (F, L, R). Scale bars, 10 µm.

**Table S1.**
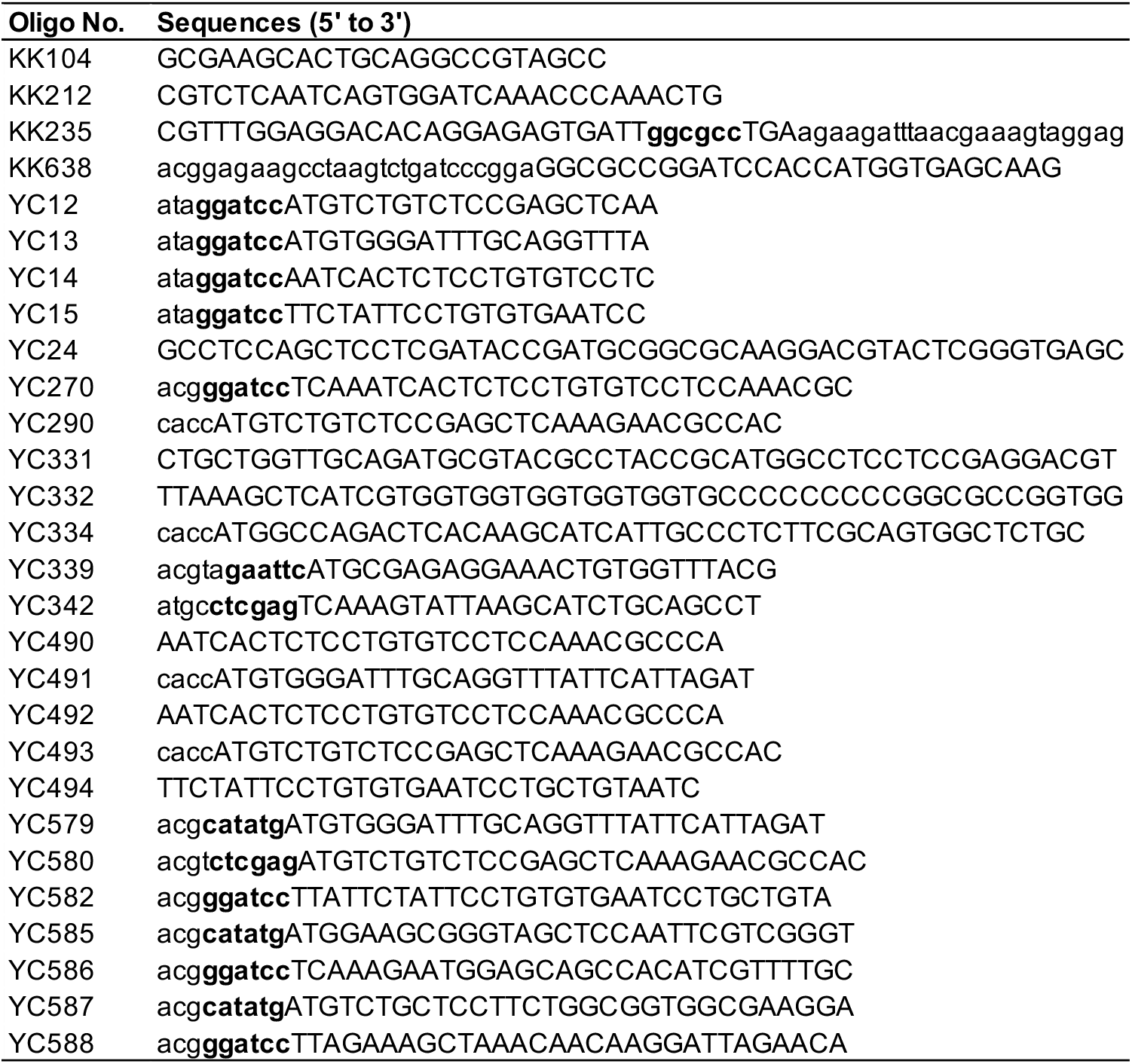
List of oligonucleotide primer sequences used in this study.

